# A Novel AuNPs-based Glucose Oxidase Mimic with Enhanced Activity and Selectivity Constructed by Molecular Imprinting and O_2_-Containing Nanoemulsion Embedding

**DOI:** 10.1101/352849

**Authors:** Lin Fan, Haoan Wu, Doudou Lou, Xizhi Zhang, Yefei Zhu, Ning Gu, Yu Zhang

**Author notes:** These authors are the corresponding authors.

## Abstract

In spite of the competitive advantages, inorganic nanoparticle mimic enzymes exhibit inherent disadvantages of limited catalytic efficiency and lacking selectivity. Here a AuNPs based mimic enzyme with significantly enhanced glucose selectivity and catalytic activity was constructed and demonstrated for the first time. Aminophenylboronic acid was employed to increase the affinity to glucose, as well as build molecular imprinted polymer shells to realize the selectivity for template molecules of glucose. Besides that, heptadecafluoro-n-octyl bromide nanoemulsion with the function of providing oxygen was introduced to gain a further improvement in catalytic activity, which successfully enhanced the catalytic efficiency (*k*_*cat*_/*K*_*m*_) up to about 270-fold. Based on the demonstrated catalytic properties, AuNPs based glucose oxidase mimics have been successfully applied in practical glucose detection of drinks and blood glucose.

## Introduction

Since Yan X and her coworkers^[1]^ brought forward the concept of enzyme nanomimics, tens of nanostructural mimic enzymes have been discovered^[2]^. Most research of mimic enzyme are focused on improving the catalytic activity, exploring the catalytic mechanism, applying to tumour diagnosis and therapy and other aspects^[3–7]^. In spite of the competitive advantages, lacking substrate-selectivity is still a significant deficiency for mimic enzymes, and there also remains a broad improvable space in catalytic activity.

Various ideas on engineering substrate-selectivity have been reported. Catalytic antibody proposed by Lerner in 1986 was one of the most influential solutions^[8]^. These catalytic immune globulins retained the characteristic of antigen-specific binding, and also exhibited enzyme-like catalytic activity^[9]^. Inspired by this work, research on catalytic antibody peaked during the 1990s, but has gradually declined in recent years due to the complicated preparation and limited catalytic efficiency. Itamar Willner^[10]^ proposed another attempt in 2015. They linked a DNA recognition sequence (aptamer) to the catalytic DNAzyme to selectively oxidize the targets of the aptamer, which leads to a 20-fold increase in catalytic activity. Kelong Fan and his coworkers^[11]^ modified Fe_3_O_4_ with histidine, which effectively enhanced the catalytic activity and affinity but still possess insufficient selectivity. Molecular imprinting technology^[12]^ is an ideal method to construct selectively binding structure, and has great applied potential in mimic enzyme with higher selectivity and affinity. Juewen Liu^[13–14]^ employed TMB and ABTS as template molecules to realize substrate-selective catalysis of Fe_3_O_4_ based mimic enzymes by molecular imprinting, which resulted in a 100-fold increase in catalytic efficiency.

As glucose (Glu) plays a crucial role in a variety of physiological activities, gold nanoparticles (AuNPs) catalyzed Glu oxidation shows important significance in both theoretical and applied research^[4,15–31]^, and has been widely applied in industrial production, scientific research and biomedical detection. On one side, the electron transfer happens between glucose oxidase (GOD) and small-sized AuNPs, which would promote the Glu oxidation catalyzed by natural enzyme^[26-27]^. On the other side, AuNPs also work as a well-behaved GOD mimic^[15,28]^. Besides that, Yihui Hu^[32]^ also designed AuNPs@MIL-101 by in situ growing AuNPs into a highly porous and thermally stable metal-organic framework, and used it as both peroxidase mimics and surface-enhanced Raman scattering substrates to realize in vitro detection of Glu and lactate. AuNPs based Glu oxidation has a great potential in biomedical application. Here, we used catalytic AuNPs to construct a molecular imprinting based substrate-selective mimic enzyme with enhanced substrate selectivity and catalytic activity. Aminophenylboronic acid (APBA) which can bind to the adjacent hydroxyls of saccharides under alkaline conditions, was conjugated to AuNPs to increase the affinity to Glu and then polymerized to engineering molecular imprinted shells. On this basis, heptadecafluoro-n-octyl bromide (PFOB) with the function of providing oxygen was introduced and further enhanced the catalytic efficiency by about 270-fold. The AuNPs based GOD mimics could be applied in practical glucose detection of drinks and blood glucose.

## Results and discussion

### Size-dependent GOD-like Activity of AuNPs

We prepared AuNPs with a diameter of 5-60 nm to study the enzyme-like activity (Figure 1abc & Supplementary Table 1). AuNPs with a diameter of 5nm and 10nm were prepared by sodium citrate-tannic acid method and AuNPs from 15nm to 60nm were prepared by kinetically controlled seeded growth synthesis^[33]^. AuNPs of different sizes were all modified with citrate and showed satisfactory water-solubility and monodispersity.

**figure 1.**
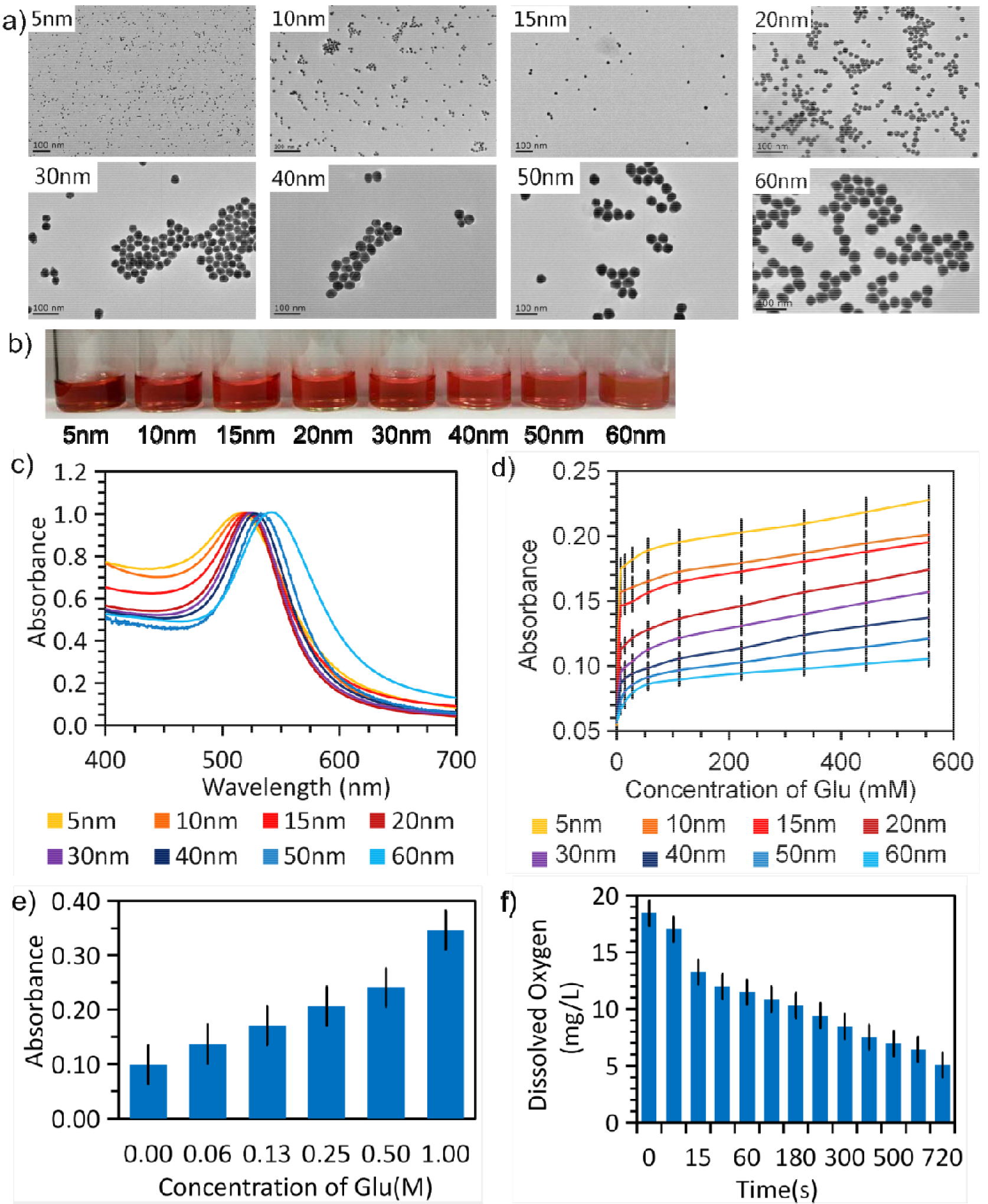
Synthesis of AuNPs from 5nm-60nm and exploration of the reaction process of AuNPs catalyzed Glu oxidation, (a) TEM images, (b) photos, (c) UV-vis absorption spectra of AuNPs. (d) Size dependency of enzyme-like activities of AuNPs for Glu oxidation, (e) The production of GIA during AuNPs catalyzed Glu oxidation, (f) The reduction of dissolved oxygen during AuNPs catalyzed Glu oxidation.

The Glu oxidation reaction catalyzed by AuNPs of different sizes was evaluated by ABTS based H_2_O_2_ detection. We removed the free tannic acid in aqueous solution of AuNPs of 5nm and 10nm by ultrafiltration before catalyzing the Glu oxidation, because that a large quality of tannic acid was needed for controlling AuNPs in the range of small size and the reductive tannic acid significantly inhibited the ABTS based colour reaction evoked by Glu oxidation^[34]^. Seen from the results, the catalytic activity of AuNPs from 5nm to 60 nm exhibited obvious downtrend with size (Figure 1d). Considering that AuNPs of 10nm were more easy to be prepared and treated compared with 5nm AuNPs, we chose AuNPs of 10nm for the follow experiment.

As mentioned above, Glu oxidation catalyzed by AuNPs could initiate ABTS coloration, which proved the production of H_2_O_2_. A production of gluconic acid (GIA) was also found through a measureable color change after reacted with diamine tetraacetic acid and ferric trichloride in sequence (Figure 1e). Besides that, a reduction of dissolved oxygen was detected by the Multi-parameter analyser during the AuNPs catalyzed Glu oxidation (Figure 1f). According to the above experimental results, it was confirmed that AuNPs induced similar Glu oxidation process with GOD^[18-19]^:

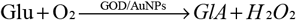

Therefore, increasing 0_2_^[31]^ involved in the reaction and improving the affinity of AuNPs for Glu could both promote the Glu oxidation. This provided theoretical support for the follow-up researches.

### Construction of MIP with Enhanced Activity and Selectivity

Although bare AuNPs has oxidase-like activity for Glu and other similar saccharides^[21,31]^, they have a significant limit of lacking selectivity and wide improvement space in catalytic efficiency. To improve the selectivity, molecular imprinting was introduced to construct mimic enzyme with enhanced catalytic activity and specific recognition of Glu (Scheme 1).

APBA was employed as recognition molecule, which could bind to the adjacent hydroxyls of saccharides under alkaline or neutral conditions (Figure 2a). APBA was adsorbed to AuNPs through both electrostatic adsorption and N-Au bonds between AuNPs and the amine groups of APBA. According to the results of enzymatic kinetics, coating with APBA successfully improved the catalytic activity of AuNPs (Figure 2b). *k*_*cat*_/*K*_*m*_ was used as a measure of enzyme efficiency^[13-14]^. According to calculation, coating with APBA lead to a smaller *K*_*m*_ and a larger *k*_*cat*_/*K*_*m*_, which implied a higher affinity to Glu and a raised catalytic efficiency (Supplementary Table 2). Compared with cholesterol (CHOL) and uric acid (UA) catalyzed by cholesterol oxidase (COD) and urate oxidase (UOD), which had similar oxidation process as Glu but without adjacent hydroxyls, coating with APBA only selectively improved the catalytic activity for Glu, while APBA-Au and bare AuNPs performed similar catalytic level for UA and CHOL, as demonstrated (Figure 2c).

**figure 2.**
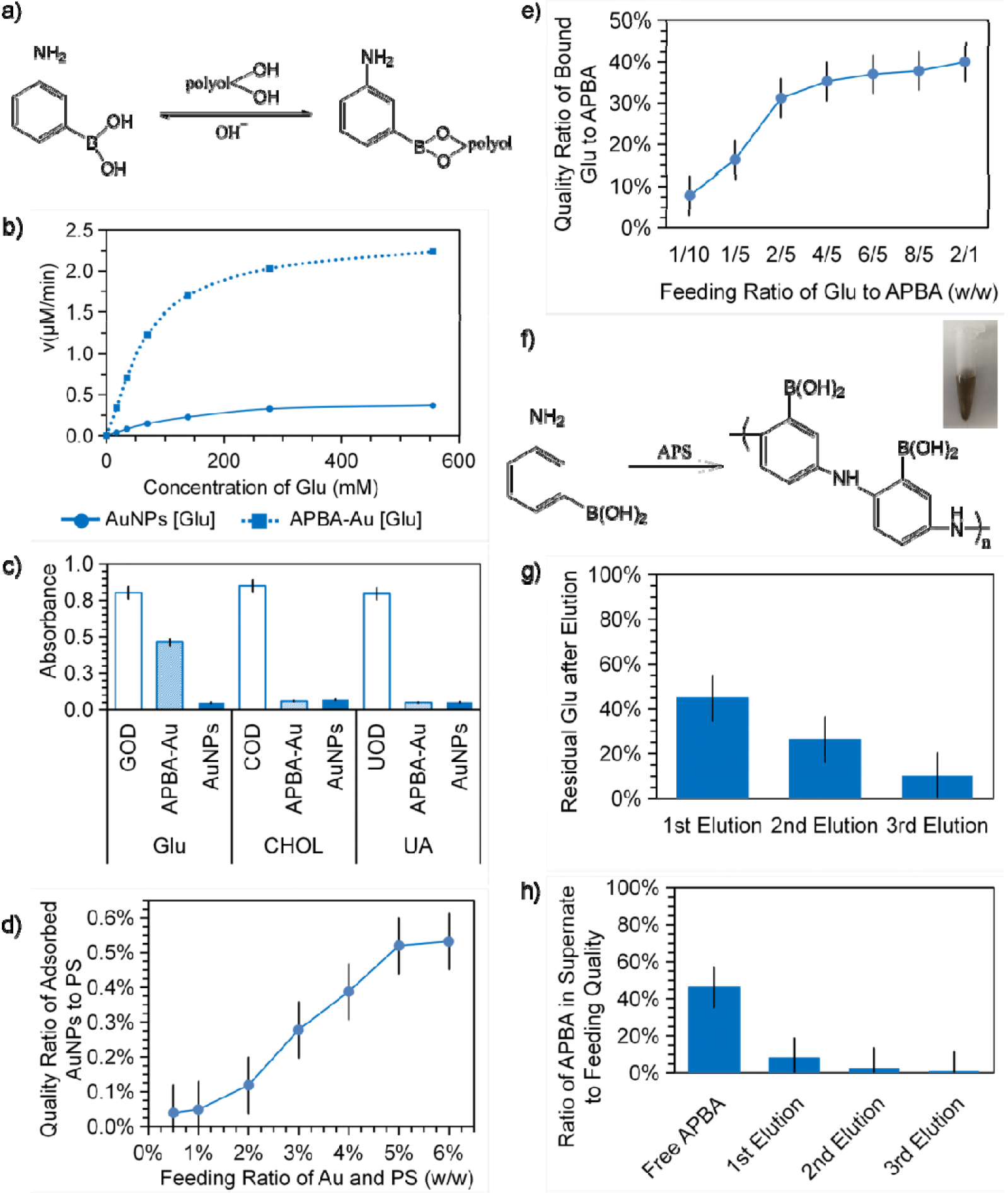
Principle and progress of the construction of MIP. (a) The reaction pathway of APBA binding to the adjacent hydroxyls of saccharide under alkaline or neutral conditions, (b) The enzymatic kinetics of AuNPs and APBA modified AuNPs catalyzed Glu oxidation, (c) Selectivity study of APBA modified AuNPs using Glu, CHOL and UA with same molarity as substrates, (d) The relationship curve between the feeding ratio and the quality ratio of adsorbed AuNPs to PS. (e) The relationship curve between the feeding ratio and the quality ratio of bound Glu to APBA. (f) APS initiated APBA polymerization reaction pathway and the photo of APBA polymers, (g) The centrifugally elution efficiency of the template molecules of Glu. (h) Ratio of APBA in supernate to feeding quality during eluting the template molecules and photos of the corresponding supernate.

APBA was coupled to the AuNPs on PS (APBA@Au-PS) (Figure 3c) and then bound with the template molecules of Glu under oxygen-free condition. The binding amount tend to equilibrium when the feeding ratio of Glu and APBA was 4/5. Under this condition, the quality ratio of bound Glu and APBA was about 35.30% and the utilization of Glu was about 44.12% (Figure 2e & Supplementary Table 3). Besides of binding to the adjacent hydroxyls, APBA could form brown polymers initiated by ammonium persulfate (APS) due to the reactive amino groups, which was employed to build the molecularly imprinted polymer shells (Figure 2f). In the presence of N.N’-methylenebisacrylamide (MBA) as cross-linking agent, polymerization took place between free APBA and those adhered on the surface of Au-PS to form a network structure (Supplementary Figure 2a). Then the polymers deposited on the surface of microspheres to form a shell of APBA around Au-PS.

**figure 3.**
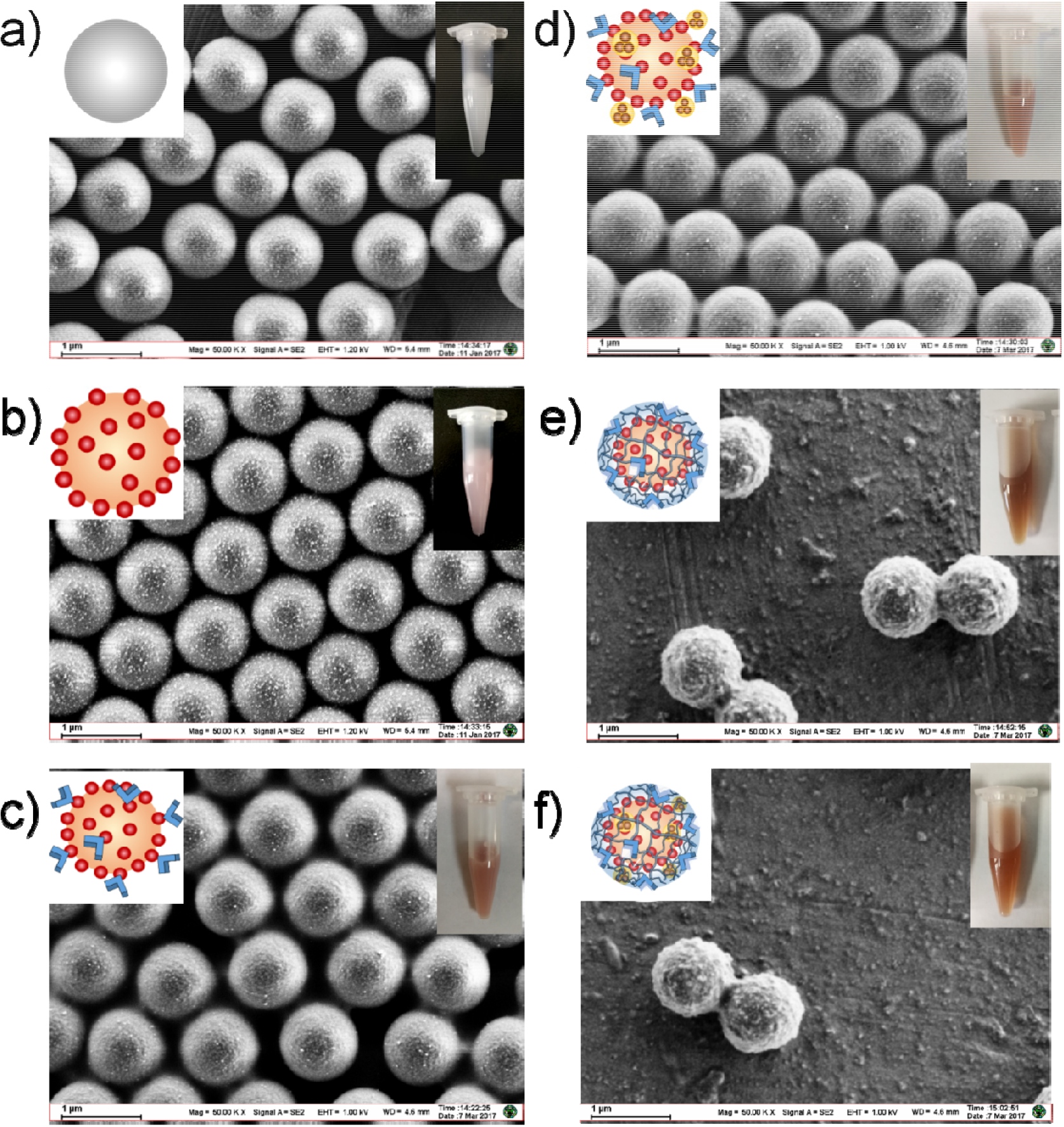
SEM images of (a) bare PS, (b) Au-PS, (c) APBA modified Au-PS, (d) APBA and PFOB modified Au-PS, (e) MIP and (f) PFOB-MIP.

As a result of that the bond between APBA and adjacent hydroxyls of Glu would break under acidic condition, the template molecules were eluted with acidic phosphate buffer (PBS) and centrifugally removed. A total of about 89.73% of the bound Glu could be washed off through three times of centrifugally elution (Figure 2g & Supplementary Table 3).

**Scheme 1.**
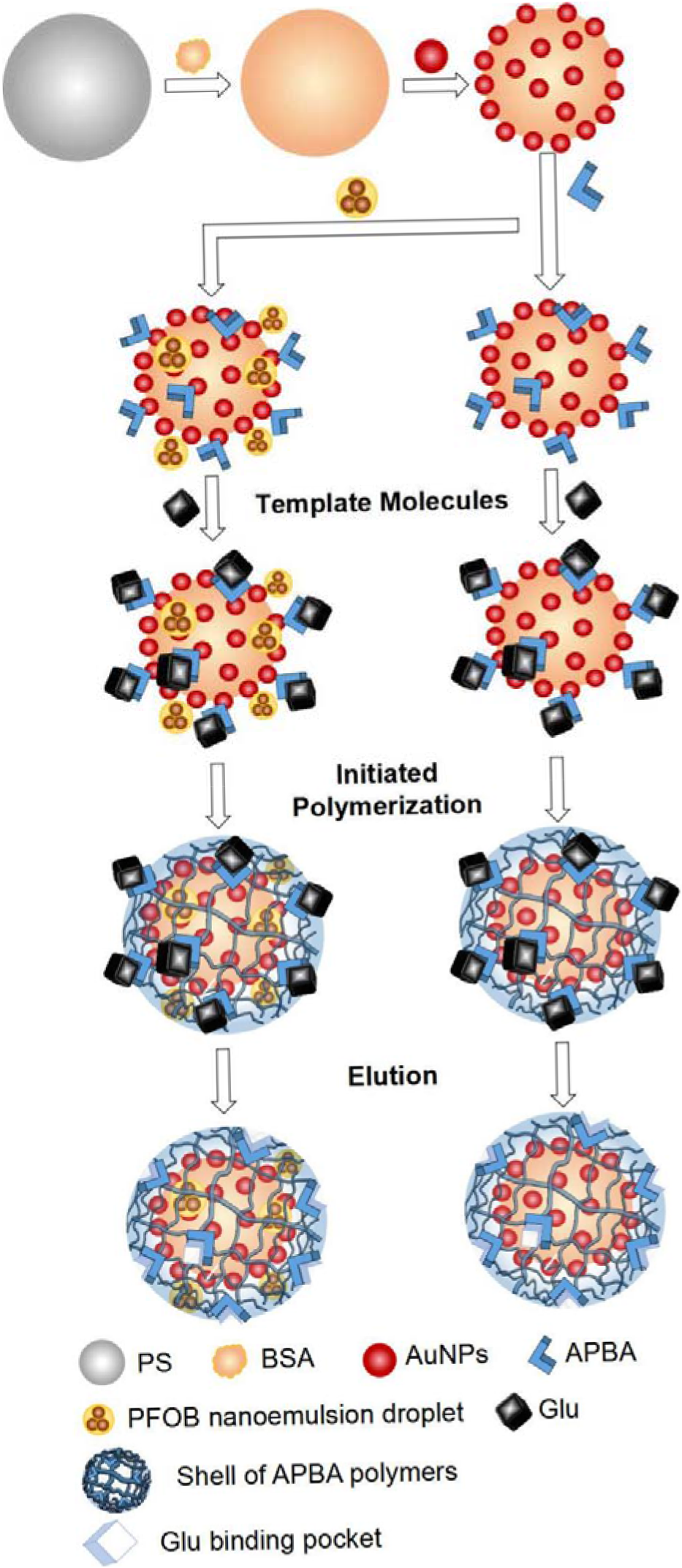
Principle of the AuNPs-based GOD mimic with enhanced activity and selectivity constructed by molecular imprinting.

MIP exceeded 500nm could be basically separated from free APBA polymers in the reactive system by centrifugation (Supplementary Figure 2). As benzene rings had characteristic peaks at 250-300nm, the centrifugally removed APBA polymers were detected by ultraviolet absorption. 46.13% of the feeding APBA were removed during the first centrifugation and little free APBA polymers were remained after three times of elution (Figure 2h & Supplementary Table 3).

After removing the template molecules, a shell with Glu binding pockets was formed on Au-PS and the rough shells of molecularly imprinted polymers (MIP) could be seen clearly under scanning electron microscope (SEM) (Figure 3e & Supplementary Figure lb). Seen from the SEM images, the APBA polymer shells were stable during centrifugally removing the template molecules and the utilization of APBA to construct MIP was about 42.99% (Figure 2h & Supplementary Table 3). Compared MIP with bare PS, the average size based on SEM increased from 0.97μm to 1.07μm and the hydrodynamic size increased from 1.05μm to 1.15μm, with good monodispersion.

To further explore the formation of APBA polymer shells, we also prepared MIP in the presence of twice amount of APBA in the reaction system. Seen from the SEM image, the shells of APBA polymers were thick and adhesive due to the excess free APBA in the system before polymerized (Supplementary Figure 3b). In an additional reaction system, the free APBA were removed from the mixture of Au-PS and APBA by centrifugation before polymerization. As a result, the polymerization could hardly take place to form the imprinted shells (Supplementary Figure 3a). These indicated that all the APBA adsorbed on the surface of Au-PS is insufficient to form polymer shell and the free APBA in reactive system contributed mainly the MIP formation.

Based on the mechanism study on AuNPs catalyzed Glu oxidation, PFOB with the function of reserving oxygen was employed to construct GOD mimics with higher catalytic activity. PFOB was encapsulated by phospholipid with amino and dispersed in water phase as nanoemulsion droplets (PL@PFOB) with a hydrodynamic size of about 26.01nm. We coupled both PL@PFOB and APBA to the surface of Au-PS at the same time (Figure 3d) and built the molecular imprinted shell with Glu binding pockets as above. The MIP with PFOB (PFOB-MIP) exhibited good dispersion (Figure 3f & Supplementary Figure lc), and the average size based on SEM and the hydrodynamic size was 1.08μm and 1.18μm respectively.

Based on the statistical results of three repeated preparations in succession, this methods was repeatable and had satisfactory standard deviations of conjugating efficiency at each steps (Supplementary Table 3). It was feasible to realize large-scale preparation. The production at gram-scale could attain similar results as small batch preparation (Supplementary Figure 3cd). And it could also be extended to construct oxidase mimics of different sizes and other saccharides with enhanced catalytic activity and substrate-selectivity.

### GOD-like Activity of MIP and Glu Detection Applications

According to the Glu-concentration dependences (Figure 4a), all the AuNPs based mimic enzyme catalyzed Glu oxidation showed linear increasing tendency in the range of experimental concentration. Compared with AuNPs, the introduction of molecular imprinting achieved a significant improvement in catalytic activity. The molecular imprinted APBA polymer shells appeared as porous structures with strong adsorbability and provided more binding sites and specific binding pockets to capture and enrich Glu around the active centers. Glu spcifically bound to the surface of AuNPs could be oxidized by activated oxygen with a higher efficiency^[36]^. Not only resulted from the increased affinity to substrate, the enhanced catalytic activity might also benefit from the effect of confinement introduced by molecular imprinted shells, which could lead to an promoted reaction rate as the demonstrated reports proposed^[37–39]^. Beyond that, it could also be seen from the enzymatic kinetics (Figure 4b) and reaction-time curve (Figure 4c) that PFOB-MIP possessed a further enhanced catalytic activity on the basis of MIP due to the providing oxygen function of PFOB. When exposed to the air, PFOB with a high solubility to O_2_ absorbed the dissolved oxygen in water until reaching a certain concentration difference. During the PFOB-MIP catalyzed Glu oxidation reaction, O_2_ in the environment was consumed and kept reducing, which broke the balance of concentration. As the concentration difference continuously changing, PFOB kept releasing O_2_ to reach new equilibrium. The MIP(removing free APBA) could not successfully form the molecular imprinted shells and thus hardly obtained improved selectivity and enhanced catalytic activity. As a result of the excessive APBA, the polymer shells around the MIP(extra APBA) was over-thick, which lead to an increase in the distance between substrate and AuNPs, as well as a decrease in catalytic activity. Note that, the catalytic activity of non-imprinted polymers (NIP) without Glu binding pockets showed no distinct improvement. As the structure optimizing, *K*_*m*_ was decreasing while V_max_ and *k*_*cat*_/*K*_*m*_ was increasing, which means stronger affinity to substrate and higher maximal velocity and catalytic efficiency (Supplementary Table 2 & Figure 4def). Compared with bare AuNPs, the catalytic efficiency of MIP and PFOB-MIP were improved by 127.05-fold and 270.92-fold, respectively (Figure 4g). Because mimic enzyme based on AuNPs had much more active centers than natural GOD, the *K*_*cat*_′ of a single GOD molecule of 10000U/mg was 4.1528×10^−21^s^−1^ while a single MIP and PFOB-MIP could reach up to 8.5284×10^−6^s^−1^ and 1.0542×l0^−5^s^−1^, respectively (Supplementary Table 4). Mal, Fru and Gal were used to explore the selectivity, and MIP exhibited higher enzymatic activity for Glu specifically (Figure 4h). We tested the reusability of PFOB-MIP for 6 cycles. PFOB-MIP was centrifugally eluted adequately and stood for several hours before the next cycle each time. PFOB-MIP kept 98.55% of the catalytic activity after 3 times of reuse. However, the catalytic activity was going down after repeatedly reused which might be caused by the reducing function of PFOB (Figure 4i).

**figure 4.**
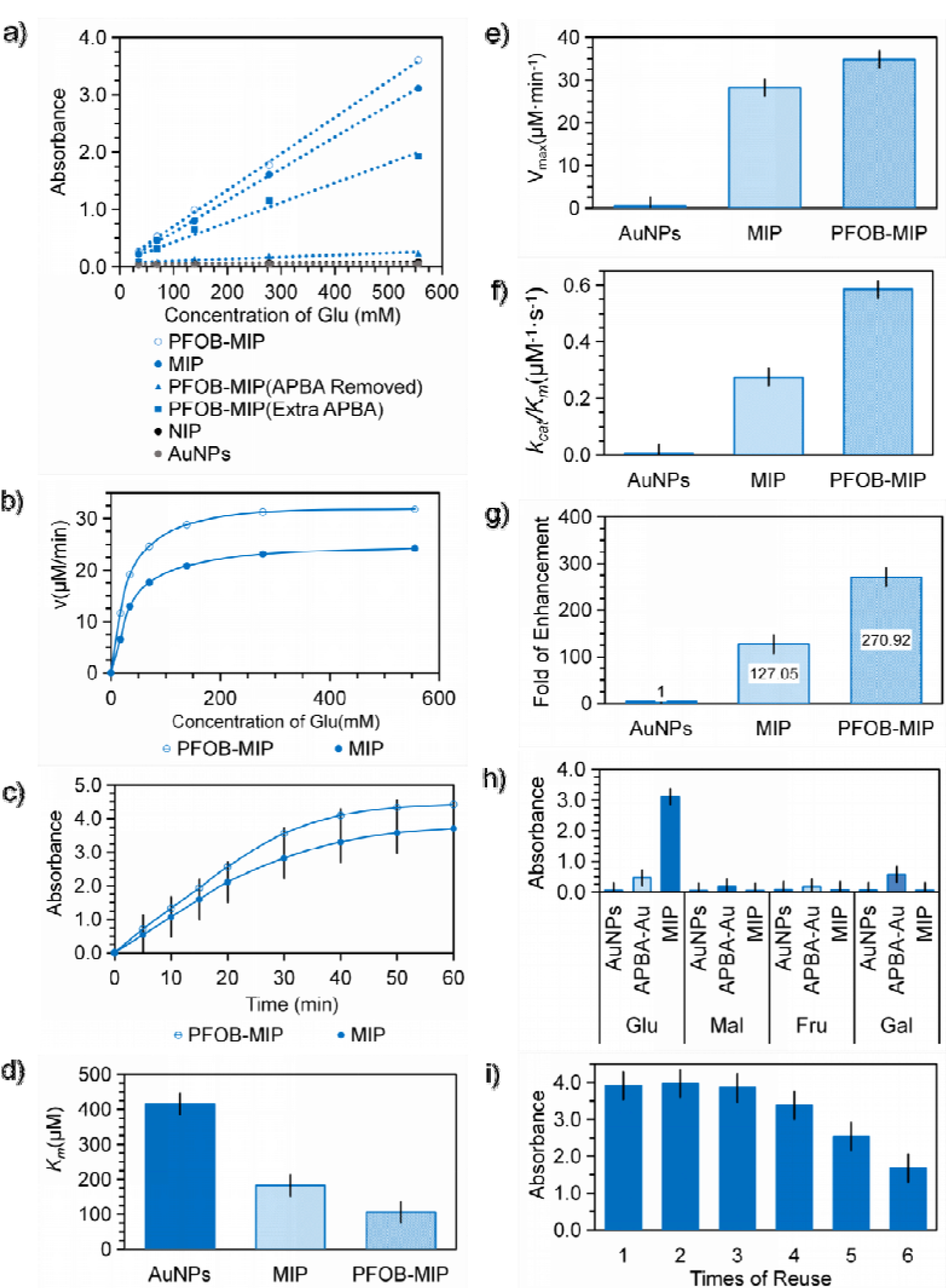
Catalytic properties of AuNPs based GOD mimics, (a) Glu concentration dependences of AuNPs based GOD mimics catalyzed Glu oxidation, (b) The enzymatic kinetics and (c) time dependences of MIP and PFOB-MIP catalyzed Glu oxidation. (d)*K*_*m*_, (e) V_max_, (f) the catalytic efficiency (*k*_*cat*_/*K*_*m*_) and (g) the fold of catalytic efficiency enhancement of AuNPs based GOD mimics catalyzed Glu oxidation, (h) Specificity study of MIP using Glu, Mal, Fru and Gal with same molarity as substrate, (i) Reusability of PFOB-MIP. Glu with a concentration of 556.56mM was catalyzed by PFOB-MIP with the same quality.

Due to extensive binding affinity of APBA for all saccharides, this method could also be extended to constructing selective mimic enzymes for other saccharides by changing the template molecules and the suitable nanoparticles.

Based on the demonstrated catalytic properties, AuNPs based GOD mimics were applied in practical detection. We used PFOB-MIP to detect the Glu in serum samples and common drinks. For the blood Glu detection, we choose serum with 2.4, 4.7,10.7,14.7, 20.6, 25.0, 30.2mM Glu as experimental samples, which basically covered the common range of blood Glu. Compared with AuNPs, PFOB-MIP showed obviously enhanced catalytic activity when detecting Glu in complex mixture (Figure 5a). Besides that, PFOB-MIP provided similar results with GOD, and exhibited good linearity with a satisfactory correlation coefficient of 0.9931 (Figure 5b & Supplementary Table 4). PFOB-MIP was also successfully used as catalyzer in food analysis. Watson Spring Water, Budweiser Beer, Coca Cola, Seven-Up, bottled Nescafe and instant Lipton Milk Tea were detected along with 555.56mM Glu solution and pure water. The concentration of Glu in different drinks were calculated by UV-vis absorption values, accounting for the results of positive (555.56mM Glu) and blank (pure water) control groups, and PFOB-MIP provided similar results without significant statistical difference with GOD (Figure 5c).

**figure 5.**
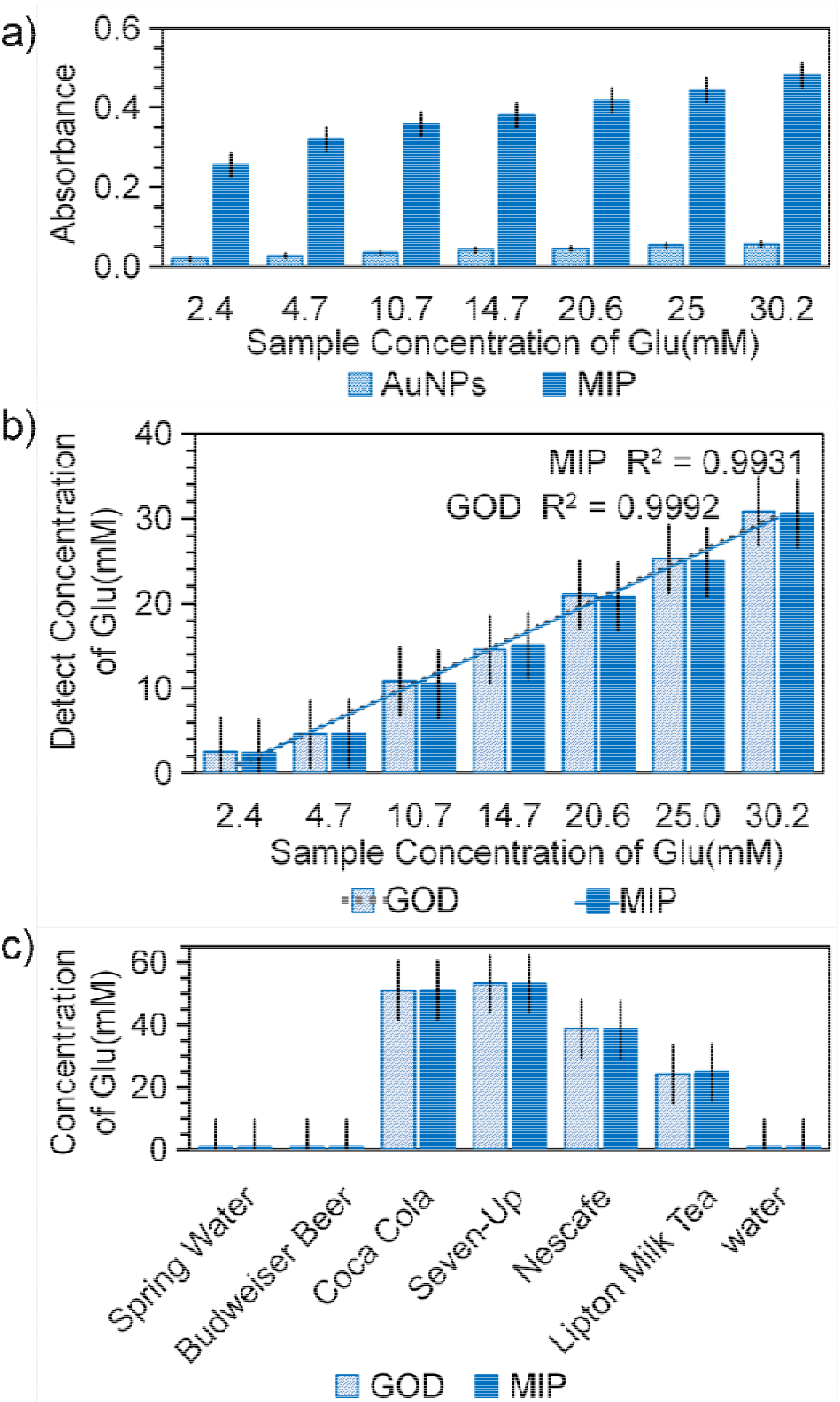
Practical Application of PFOB-MIP. (a) The comparison of detection results of blood Glu using AuNPs and PFOB-MIP as catalyzer, and (b) the calculation results of Glu content detected by GOD and PFOB-MIP. (c) The Glu content detection results of Glu in common drinks (Watson Spring Water, Budweiser Beer, Coca Cola, Seven-Up, bottled Nescafe, instant Lipton Milk Tea, positive control (555.56mM Glu) and blank control (pure water) using GOD and PFOB-MIP as catalyzer.

## Experimental

### Preparation of APBA-Au

Sodium citrate-tannic acid method were used for preparing gold nanoparticles with a diameter of 5nm and l0nm. Chloroauricacid (lml, 0.0lg/ml, Aladdin Industrial Corporation), potassium carbonate (100μl for 5nm and 25μl for 10nm, 0.1M), trisodium citrate (4ml, 0.01g/ml) and tannic Acid (700μl for 5nm and 100μl for l0nm, 0.0lg/ml) were mixed in a system with a volume of 100ml and stirred at 60 ⍰ for 10 min, and then cooled to room temperature. Gold nanoparticles from 15nm to 60nm were prepared by kinetically controlled seeded growth synthesis^[33]^ and diluted to 0.05mg/ml. Unless stated, AuNPs with a diameter of 10nm were used for construction of selective mimic enzyme.

APBA-Au were prepared by electrostatic attraction. APBA (50μl, 2mg/ml, Aladdin Industrial Corporation) was mixed with AuNPs with a diameter of l0nm (lml, 0.05mg/ml) and incubated for 2h. The mixture was then centrifuged to remove unbound APBA at 12000 rpm for 40 min at 4⍰.

### Preparation of MIP and PFOB-MIP

PS were used as supporters. PS of 1000nm (200μl, 50mg/ml, Nanomicro Tech) were incubated with BSA (10mg, YEASEN) overnight and centrifugally washed throughly. Deposition was resuspended with an aqueous solution containing AuNPs with a diameter of 10nm (10ml, 0.05mg/ml) and incubated for 2h.The mixture containing Au-PS was then centrifuged to remove unbound AuNPs^[40]^. The amount of Au in Au-PS was calculated by UV absorption value of the feeding AuNPs and the unbound AuNPs. PS of other sizes were prepared in the same way and only PS of 1000nm were used for follow-up researches.

PFOB was emulsified with surfactant to dissolve in water phase. Phospholipid with amino (PL-NH_2_) (50mg, Nanosoft Biotechnology LLC) and chloroform (2ml) were mixed in a system with a volume of 20ml and stirred intensely at 100 ⍰, and PFOB (500μl, Alligator Reagent) was added dropwise. The mixture containing phospholipid emulsified PFOB (PL@PFOB) was then cooled to room temperature and ultrasonically dispersed.

MIP were prepared under oxygen-free condition. Under N_2_ PS-Au were incubated with APBA (5mg), Glu (4mg) and MBA (50mg, Biosharp) for 2h in order, then mixed with APS (0.8ml, 22.82mg/ml) and PBS (10ml, 0.02M, pH9) for 10min, and then heated at 60 ⍰ overnight. The MIP were washed throughly with PBS (0.02M, pH5) to remove the template molecules of Glu as well as the free APBA polymers. The centrifugally removed APBA polymers were detected by UV-vis. Then the MIP were centrifugally collected and dispersed in distilled water.

The production of MIP was scaled up to 1g. PS of 800nm (10ml, 100mg/ml, Nanoeast Tech) were employed to promoting the utilization of AuNPs. PS were incubated with BSA (1g, YEASEN) overnight and centrifugally washed throughly. Deposition was resuspended with an aqueous solution containing AuNPs with a diameter of 10nm (100ml, 0.05mg/ml) and incubated for 2h.The mixture containing Au-PS was then centrifuged to remove unbound AuNPs and dissolved in 20ml of distilled water. Under oxygen-free condition PS-Au were incubated with APBA (500mg), Glu (400mg) and MBA (5g, Biosharp) for 2h in order, then mixed with APS (40ml, 0.5g/ml) dissolved in PBS (pH9) for 10min, and then heated at 60 ⍰ overnight. The MIP were centrifugally collected after washed with PBS (0.02M, pH5) to remove the template molecules of Glu and dispersed in distilled water.

The PFOB-MIP were prepared in the same way except that PL-NH_2_@PFOB (5ml, 2.5%, v/v) was added along with APBA. NIP were also prepared in the same way except that no template molecules of Glu was added.

To explore the formation regularity of APBA shells, MIP(removing free APBA) and MIP(extra APBA) with different contents of APBA was prepared. The free APBA in MIP(removing free APBA) were removed by centrifugation before incubated with MBA. Extra 5mg APBA was added into MIP(extra APBA) before mixed with APS. The rest of the preparation steps were in the same way as the MIP.

The above particles were evaluated by transmission electron microscopy (TEM) (JEOL, JEM-2100), scanning electron microscope (SEM) (Ultra Plus, Zeiss), Energy Dispersive X-Ray Spectroscopy (EDX) (Ultra Plus, Zeiss), ultraviolet visible spectroscopy (Uv-vis) (Shimadzu Scientific Instruments, UV-3600) and dynamic light scattering (DLS) (Brook haven, Zeta Plus).

### Activity Assays

The activity assays were performed in 96-well plates.
Glu of 555.56, 444.45, 333.34, 222.22, 111.11, 55.56, 27.78, 13.89, 6.94 and 0mM was dissolved in PBS (0.02M, pH7.4) and used as substrates for exploring the catalytic activity of AuNPs with different sizes. The substrates (50μl) was mixed with horse radish peroxidase (HRP) (1μl, 1mg/ml) and ABTS (10μl, 10mg/ml), then incubated with AuNPs (200μl, 26.5μg/ml) at 25⍰. The absorbance at 405nm was measured by Multiscan Spectrum (Tecan M200) and the reactions catalyzed by AuNPs with different sizes were measured at the same time before the one with highest reaction rate over range. Glu of 555.56, 277.78, 138.89, 70.00, 34.72, 17.36 and 0mM was dissolved in PBS (0.02M, pH7.4) and used as substrates for exploring the catalytic activity of selective enzyme mimics. Glu of 555.56mM was chosen to explore the time dependency and reusability. The substrates (50μl) was mixed with horse radish peroxidase (HRP) (1μl, 1mg/ml) and ABTS (10μl, 10mg/ml), then incubated with GOD (200μl, 5U/ml, Aladdin 10000U), AuNPs (200μl, 26.5μg/ml), APBA-Au (200μl, containing 26.5μg/ml of AuNPs) or imprinted polymers (200μl, containing 26.5μg/ml of AuNPs) at 25⍰. The absorbance at 405nm was measured by Multiscan Spectrum (Tecan M200) and the reactions catalyzed by different enzymes were measured at the same time before the one with highest reaction rate over range.

555.56mM of CHOL, UA, Mal, Fru and Gal was used as substrate to explore the selectivity of APBA-Au and MIP. The oxidation of different substrates were catalyzed by AuNPs based mimic enzymes with same molarity of AuNPs and natural enzymes with same active unit. The rest of the operation steps were in the same way as the Glu.

V_max_, *K*_*m*_ and *k*_*cat*_ were obtained by fitting the data with the Beer-Lambert Law: A=lg(1/T)=Kbc (K=36000 M^−1^cm^−1^for ABTS and b is the pathlength of 0.81 cm), and Michaelis-Menten equation: V_0_=V_max_[S]/(*K*_*m*_+[S]) and *k*_*cat*_=V_max_/[S].

The detection of Glu in blood and drinks was performed in the similar way. For the blood Glu, serum samples with 2.4, 4.7, 10.7, 14.7, 20.6, 25.0, 30.2mM Glu were adopted as substrate (the blood Glu value was given by Omron glucometer) and catalyzed by AuNPs, MIP and GOD. For the drinks, Watson Spring Water, Budweiser Beer, Coca Cola, Seven-Up, bottled Nescafe, instant Lipton Milk Tea were used as substrate and pure water was used as negative control. The drinks were catalyzed by MIP and GOD.

### GIA Detection

Glu of 1M was dissolve in distilled water and two-times step diluted to 5 concentrations along with an extra blank group with no Glu. The substrate (20μl) was incubated with AuNPs (40μl, 0.05mg/ml) or distilled water (40μl) at 37⍰ for 60mim. Then the mixture was mixed with ethylene diamine tetraacetic acid (100μl, 5mM), triethylamine (100μl, 0.15M), ammonium hydroxide (10μl, 3M). 15min later, concentrated hydrochloric acid (50μl), ferric trichloride (50μl, 0.1M), trichloroacetic acid (50μl, 0.25M) was added into the mixture and the absorbance at 700nm was measured at 5 min.

### Dissolved oxygen

Glu (10ml, 200mM) was kept in a test tube and mixed with AuNPs (10ml, 0.05mg/ml). A Multi-parameter controller (DZS-708, REX) was used to monitor the change of dissolved oxygen in the reaction system.

## Conclusions

In summary, a AuNPs based mimic enzyme with selectivity and 270-fold increased catalytic efficiency was constructed by molecular imprinting technology. The molecular imprinted shells with specific binding pockets to recognize, capture and enrich Glu were polymerized by adjacent hydroxyls-bindable APBA, which also lead to an enhanced catalytic activity due to the increased affinity to substrate. The porous molecular imprinted structure might also introduce the advantages of confinement effect. Beyond that, PFOB nanoemulsion with high solubility to O_2_ was embedded around the active centers and served as an oxygen donor, which provided a further improvement in the catalytic activity. This method demonstrated to be satisfactorily repeatable was feasible to realize large-scale preparation and extended to constructing selective mimic enzymes for other saccharides. These AuNPs based GOD mimics were successfully applied in practical Glu detection. Considering the advantage in aspect of cost and stability, enzyme-mimicking nanomaterials with an increased selectivity seemed to have great potential in biotechnological application.

## Acknowledgements

This work was supported by the National Key Research and Development Program of China [No. 2017YFA0205502]; National Natural Science Foundation of China [No. 81571806, 81671820]; the Science and Technology Support Project of Jiangsu Province [No. BE2017763]; the Jiangsu Provincial Special Program of Medical Science [No. BL2013029]; and the Fundamental Research Funds for the Central Universities.

## Notes and references

1 Lizeng Gao, Jie Zhuang, Leng Nie, Jinbin Zhang, Yu Zhang, Ning Gu, Taihong Wang, Jing Feng, Dongling Yang, Sarah Perrett & Xiyun Yan. Intrinsic peroxidase-like activity of ferromagnetic nanoparticles[J]. Nature Nanotechnology, 2007, 2(9): 577–583.

2 Wang X, Hu Y, Wei H. Nanozymes in bionanotechnology: from sensing to therapeutics and beyond[J]. Inorganic Chemistry Frontiers, 2016, 3(1): 41–60.

3 Liu B, Liu J. Surface modification of nanozymes[J]. Nano Research, 2017,10(4): 1–24.

4 Wei H, Wang E. Nanomaterials with enzyme-like characteristics (nanozymes): next-generation artificial enzymes.[J]. Chemical Society Reviews, 2013, 42(14):6060–6093.

5 Kaltenbach M, Tokuriki N. Dynamics and constraints of enzyme evolution.[J]. Journal of Experimental Zoology Part B Molecular & Developmental Evolution, 2014, 322(7): 468–487.

6 Fan L, Tian Y, Yin R, Lou D, Zhang X, Wang M, Ma M, Luo S, Li S, Gu N, Zhang Y. Enzyme catalysis enhanced dark-field imaging as a novel immunohistochemical method[J]. Nanoscale, 2016, 8(16): 8553–8558.

7 Fan L, Lou D, Zhang Y, Ning G. Rituximab-Au nanoprobes for simultaneous dark-field imaging and DAB staining of CD20 over-expressed on Raji cells[J]. Analyst, 2014, 139(22):5660–5663.

8 Tramontano A, Janda K D, Lerner R A. Catalytic antibodies.[J]. Science, 1986, 234(4783):1566–1570.

9 Lerner R A, Benkovic S J, Schultz P G. At the crossroads of chemistry and immunology: catalytic antibodies[J]. Science, 1991, 252(5006): 659–67.

10 Eyal Golub, H. Bauke Albada, Wei-Ching Liao, Yonatan Biniuri, and Itamar Willner. Nucleoapzymes: Hemin/G-Quadruplex DNAzyme–Aptamer Binding Site Conjugates with Superior Enzyme-like Catalytic Functions[J]. Journal of the American Chemical Society, 2016,138(1): 164–172.

11 Fan K, Wang H, Xi J, Liu Q, Meng X, Duan D, Gao L, Yan X. Optimization of Fe304 nanozyme activity via single amino acid modification mimicking an enzyme active site [J]. Chemical Communications, 2016, 53(2): 424–427.

12 Wulff G, Sarhan A, Zabrocki K. Enzyme-analogue built polymers and their use for the resolution of racemates[J]. Tetrahedron Letters, 1973,14(44): 4329–4332.

13 Zhang Z, Liu B, Liu J. Molecular Imprinting for Substrate Selectivity and Enhanced Activity of Enzyme Mimics[J]. Small, 2016, 13: 1602730.

14 Zhang Z, Liu B, Liu J. Molecular Imprinting on Inorganic Nanozymes for Hundred-fold Enzyme Specificity[J]. Journal of the American Chemical Society, 2017, 139(15): 5412–5419.

15 Comotti Ml, Della Pina C, Matarrese R, Rossi M. The catalytic activity of “naked” gold particles[J]. Angewandte Chemie, 2004,43(43): 5812–5815.

16 He Wl, Zhou YT, Wamer WG, Hu X, Wu X, Zheng Z, Boudreau MD, Yin JJ. Intrinsic catalytic activity of Au nanoparticles with respect to hydrogen peroxide decomposition and superoxide scavenging[J]. Biomaterials, 2013, 34(3):765–773.

17 Wang J. Electrochemical glucose biosensors[J]. Chemical Reviews, 2017, 108(2): 814–825.

18 Paolo Beltramea, Massimiliano Comottib, Cristina Della Pinab, Michele Rossi. Aerobic oxidation of glucose: II. Catalysis by colloidal gold [J]. Applied Catalysis A General, 2006, 297(1): 1–7.

19 M Comotti, C Della Pina, E Falletta, M Rossi. Aerobic Oxidation of Glucose with Gold Catalyst: Hydrogen Peroxide as Intermediate and Reagent[J]. Advanced Synthesis & Catalysis, 2010, 348(3): 313–316.

20 Ansari S A, Husain Q. Potential applications of enzymes immobilized on/in nano materials: A review.[J]. Biotechnology Advances, 2012, 30(3): 512–523.

21 Önal Y, Schimpf S, Claus P. Structure sensitivity and kinetics of d-glucose oxidation to d-gluconic acid over carbon-supported gold catalysts[J]. Journal of Catalysis, 2004, 223(1): 122–133.

22 Sari Rautiainen, Petra Lehtinen, Marko Vehkämki, Klaus Niemelä, Marianna Kemell, Mikko Heikkilä, TimoRepo. Microwave-assisted base-free oxidation of glucose on gold nanoparticle catalysts[J]. Catalysis Communications, 2016, 74: 115–118.

23 Massimiliano Comotti, Cristina Della Pina, Roberto Matarrese, Michele Rossi, Attilio Siani. Oxidation of alcohols and sugars using Au/C catalysts: Part 2. Sugars[J]. Applied Catalysis A General, 2005, 291(1-2): 204–209.

24 Hashmi ASK, Hutchings G J. Gold catalysis[J]. Angewandte Chemie, 2006, 45(47): 7896–7936.

25 Biella S, Prati L, Rossi M. Selective Oxidation of D-Glucose on Gold Catalyst[J]. Journal of Catalysis, 2002, 206(2): 242–247.

26 Xiao-yan Chen, Jin-ru Li, Xing-chang Li, Long Jiang. A New Step to the Mechanism of the Enhancement Effect of Gold Nanoparticles on Glucose Oxidase[J]. Biochemical & Biophysical Research Communications, 1998, 245(2): 352–355.

27 Z.J. Chen, X.M. Ou, F.Q. Tang, L. Jiang. Effect of nanometer particles on the adsorbability and enzymatic activity of glucose oxidase[J]. Colloids & Surfaces B Biointerfaces, 1996, 7(3): 173–179.

28 PC Della, E Falletta, L Prati, M Rossi. Selective oxidation using gold.[J]. Chemical Society Reviews, 2008, 37(9): 2077–2095.

29 Mirescu A, Prüße U. Selective glucose oxidation on gold colloids[J]. Catalysis Communications, 2006, 7(1): 11–17.

30 Gang Chang, Honghui Shu, Kai Jia, Munetaka Oyamab, Xiong Liu, Yunbin He. Gold nanoparticles directly modified glassy carbon electrode for non-enzymatic detection of glucose[J]. Applied Surface Science, 2014, 288(1): 524–529.

31 Robert Wojcieszak, Iolanda M.Cuccoviab, Márcia A.Silvab, Liane M.Rossi. Selective oxidation of glucose to glucuronic acid by cesium-promoted gold nanoparticle catalyst[J]. Journal of Molecular Catalysis A Chemical, 2016, 422: 35–42.

32 Hu Y, Cheng H, Zhao X, Wu J, Muhammad F, Lin S, He J, Zhou L, Zhang C, Deng Y, Wang P, Zhou Z, Nie S, Wei H. Surface-Enhanced Raman Scattering Active Gold Nanoparticles with Enzyme-Mimicking Activities for Measuring Glucose and Lactate in Living Tissues.[J]. Acs Nano, 2017, 11(6): 5558–5566.

33 Bastús N G, Comenge J, Puntes V. KineticaIly controlled seeded growth synthesis of citrate-stabilized gold nanoparticles of up to 200 nm: size focusing versus Ostwald ripening.[J]. Langmuir the Acs Journal of Surfaces & Colloids, 2011, 27(17):11098–11105.

34 Je-Min Choi, Jin Han, Byoung-Seok Yoon, Jae-Hwan Chung, Dong-Bum Shin, Sang-Kyou Lee, Jae-Kwan Hwang, Ryung Ryang. Antioxidant Properties of Tannic Acid and its Inhibitory Effects of Paraquat-Induced Oxidative Stress in Mice[J]. Food Science & Biotechnology, 2006,15(5):728–734.

35 Kumar A, Mandal S, Selvakannan P R, Pasricha R, AB Mandale, Sastry M. Investigation into the Interaction between Surface-Bound Alkylamines and Gold Nanoparticles[J]. Langmuir the Acs Journal of Surfaces & Colloids, 2003, 19(15):6277–6282.

36 Shen X, Liu W, Gao X, Lu Z, Wu X and Gao X. Mechanisms of Oxidase and Superoxide Dismutation-like Activities of Gold, Silver, Platinum, and Palladium, and Their Alloys: A General Way to the Activation of Molecular Oxygen[J]. Journal of the American Chemical Society, 2015,137(50):15882–15891.

37 X Wang, Cao W, L Qin, T Lin, W Chen, S Lin, J Yao, X Zhao, M Zhou, C Hang and H Wei. Boosting the Peroxidase-Like Activity of Nanostructured Nickel by Inducing Its 3+ Oxidation State in LaNiO3 Perovskite and Its Application for Biomedical Assays.[J]. Theranostics, 2017, 7(8):2277–2286.

38 Pan X, Bao X. The effects of confinement inside carbon nanotubes on catalysis.[J]. Accounts of Chemical Research, 2011, 44(8):553–562.

39 Yin Y, Chen M, Zhou S and Wu L. A general and feasible method for the fabrication of functional nanoparticles in mesoporous silica hollow composite spheres[J]. Journal of Materials Chemistry, 2012, 22(22):11245–11251.

40 Zhang X Z, Zhou Y, Zhang W, Zhang Y, Gu N. Polystyrene@Au@prussian blue nanocomposites with enzyme-like activity and their application in glucose detection[J]. Colloids & Surfaces A Physicochemical & Engineering Aspects, 2016, 490:291–299.

